# A sequence-based proactive intelligence for influenza antigenic profiling improves vaccine strain selection

**DOI:** 10.64898/2026.04.18.719333

**Authors:** Yihao Chen, Ying Xu, Yanhui Cheng, Xianzhi Qi, Tian Bai, Jiaying Yang, Huanle Luo, Xiangjun Du, Lin Zhu, Lei Yang, Mang Shi, Dayan Wang, Zhaorong Li, Yuelong Shu

## Abstract

Seasonal influenza viruses accumulate antigenic changes, eroding population immunity and necessitating recurrent vaccine updates. Hemagglutination inhibition (HI) assays are the standard for measuring antigenic relationships between circulating and vaccine strains; however, their limited throughput constrains the scale and timeliness of surveillance. Here, we present fluProfiler, a foundation-model-based framework that learns a stable mapping from viral sequences to antigenic space and uses this representation to support influenza antigenic prediction, vaccine strain evaluation, and diversity-driven sampling. fluAgPredictor aligns hemagglutinin (HA) and neuraminidase (NA) sequence representations with HI-derived antigenicity, enabling accurate and consistent inference of pairwise antigenic distances across surveillance-aligned evaluation settings. Without prior annotation of antigenic sites, it identifies key residues in immunodominant epitopes and reveals the cooperative contributions of HA and NA to antigenic variation. Building on this antigenic-space representation, fluVacSelector provides antigenic coverage scores that are concordant with World Health Organization (WHO) vaccine recommendations while also flagging potential candidates that may offer broader coverage ahead of formal consultations. fluAgEnhancer further leverages the same representation to prioritize antigenically informative and diverse strains for experimental characterization, achieving comparable predictive accuracy with approximately 25% fewer HI measurements than random sampling. Together, these modules provide a high-throughput and interpretable complement to HI testing, converting routine genomic surveillance into a more proactive, data-driven support system for antigenic monitoring and vaccine strain selection.

## Introduction

Seasonal influenza viruses continually accumulate mutations in their surface glycoproteins, hemagglutinin (HA) and neuraminidase (NA), gradually eroding the protection conferred by prior infection or vaccination^1^. To sustain vaccine effectiveness, the WHO Global Influenza Surveillance and Response System (GISRS) monitors circulating strains and periodically updates vaccine composition when substantial antigenic drift is detected^2^.

Central to this process is the hemagglutination inhibition (HI) assay, which remains the standard for measuring antigenic relationships between vaccine strains and circulating viruses. However, HI testing is labor-intensive and is limited in throughput. Only a limited number of strains can be phenotyped each season. Consequently, GISRS must infer the cross-protection between vaccine and circulating viruses from sparse HI data, potentially delaying the identification of emerging antigenic clusters and underestimating the erosion of vaccine efficacy.

To address these limitations, computational methods have been developed to use sequence and HI data to support more systematic antigenic surveillance^3-7^. Early work, such as antigenic cartography, embedded viral strains and antisera into a continuous antigenic map by minimizing the error between antigenic and Euclidean distances^1^. Subsequent models, such as Nextflu^5^, predicted antigenic distances by decomposing them into serum potency, viral activity, and genetic differences represented by phylogenetic distances or specific mutations. More recently, season-by-season frameworks have aligned model training with the surveillance timeline to enable the forecasting of future influenza seasons^4^. Together, these studies established important computational frameworks for antigenic visualization, prediction, and prospective surveillance. Nevertheless, most were not designed to learn an antigenicity-focused sequence representation that remains transferable under continual drift.

Continual mutation reshapes the antigenic patterns of influenza over time. As the antigenic effects of substitutions depend on the genetic context and the surrounding fitness landscape evolves^8-10^, sequence–serology mappings inferred from historical data may not remain fully stable when novel combinations of substitutions arise in later viruses. In effect, antigenic prediction becomes akin to aiming at a moving target: the model extrapolates from yesterday’s rules while the target continues to shift.

Large-scale pretraining enables transformer foundation models to learn contextual protein embeddings that capture both local motif and long-range dependencies^11-13^. Because this representation space is structured by evolutionary relationships, it can provide a useful basis for transfer across influenza subtypes and for reasoning about previously unseen mutation combinations^14-16^. In practice, pretrained sequence representations can be aligned with sparse phenotypic supervision to improve data efficiency^4,17^, with potential advantages for antigenic prediction under continual drift. However, general-purpose embedding spaces are not antigenicity optimized. Only a subset of sequence-derived signals is mechanistically relevant to HI^18-20^, and without explicit task alignment models may mistakenly attribute antigenic meaning to background variation (for example, stability constraints or replication adaptation)^9,20-22^, potentially yielding strong retrospective fits yet weaker generalization and calibration on new strains, future seasons, or novel mutational constellations^4,23^.

Here we move beyond predicting pairwise HI-derived distances to learn a stable mapping from sequence to antigenic space. Starting from a pretrained viral foundation model, we add a task-aligned attention-based extractor that reweights general protein embeddings toward antigenically informative features, improving robustness under continual drift while enabling residue-level interpretability. Building on this mapping, we present fluProfiler, comprising fluAgPredictor for sequence-based HI distance prediction, fluVacSelector for prospective vaccine-coverage scoring and candidate ranking, and fluAgEnhancer for diversity-driven active learning to prioritize HI assays that anchor sparsely sampled regions of antigenic space—together converting routine genomic monitoring into actionable support for vaccine strain selection.

## Results

### Experimental design

We seek a transferable sequence-to-antigenic-space mapping that remains stable under drift, beyond retrospective regression of pairwise HI readouts. In operational surveillance, HI evidence is intrinsically sparse for three reasons. First, HI assays do not cover all serum–virus pairs, leaving systematic gaps in the HI table. Second, only a fraction of circulating strains can be reliably phenotyped, owing to isolation and throughput constraints. Third, phenotyped viruses are typically tested against a small serum panel, truncating observed cross-reactivity and narrowing inferred cross-protection boundaries^24-26^. Crucially, strain selection occurs within a fixed decision window, making the task inherently prospective rather than retrospective under complete information^3,4,27^.

fluAgPredictor (Fig. 1B) is designed to operate under the sparsity and latency that characterize surveillance HI data. Therefore, the evaluation was framed around four surveillance-realistic scenarios that progressively increase in stringency and map directly onto operational constraints. First, sparse HI matrices were completed under titer missingness. Second, extrapolation was assessed for viruses lacking HI measurements under strain missingness. Third, generalization to previously unseen antisera was evaluated under serum missingness. Finally, strictly timeline-aligned, cross-season prospective prediction was tested to reflect the fixed decision window of vaccine strain selection (Fig. 2A).

**Figure 1.**
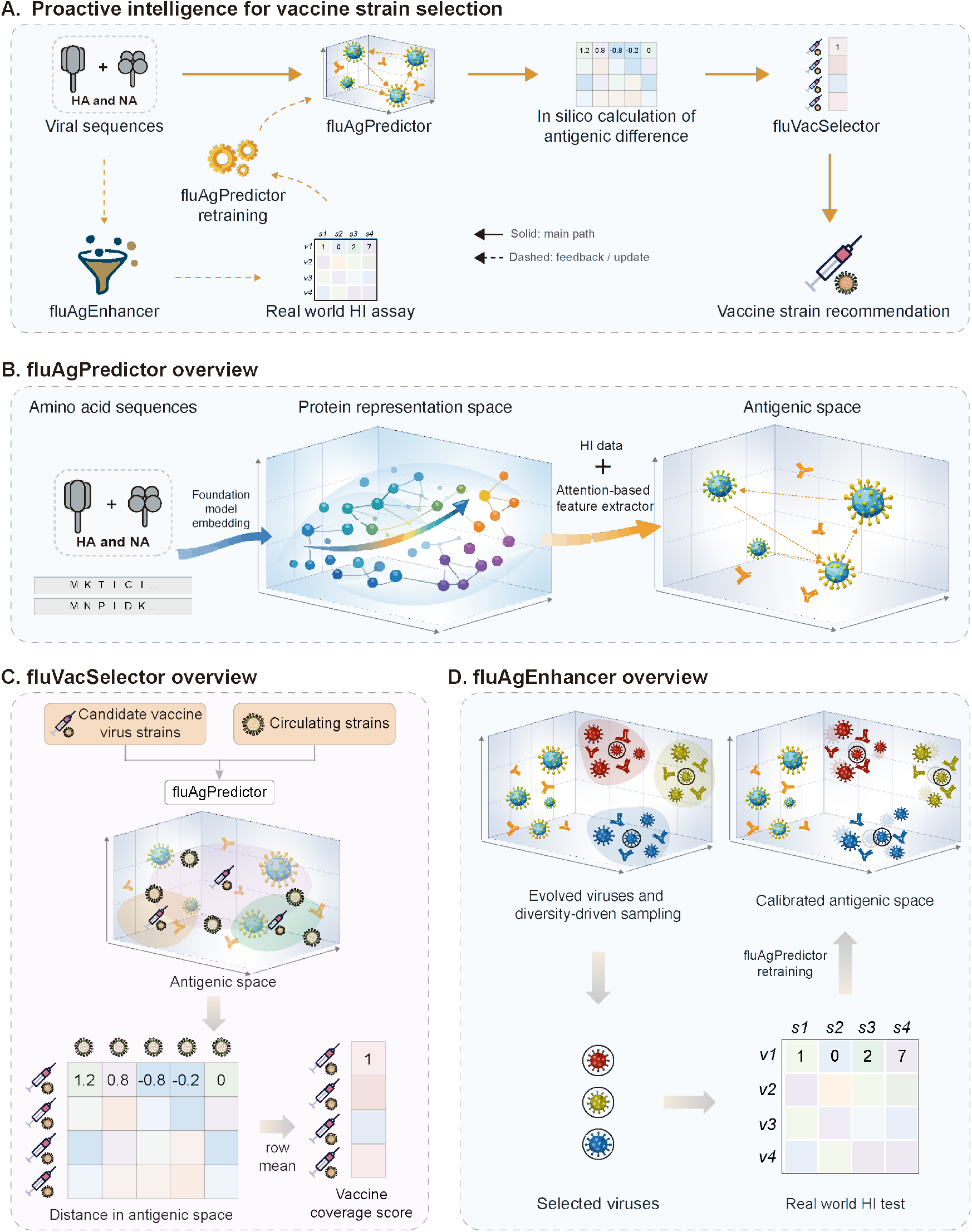
fluProfiler framework for antigenic prediction, vaccine recommendation and diversity-driven sampling. A, Overview of fluProfiler. Viral HA and NA sequences are used by fluAgPredictor to infer pairwise antigenic distances, by fluVacSelector to rank candidate vaccine strains by predicted coverage, and by fluAgEnhancer to select informative viruses for HI testing and iterative model updating. Solid arrows indicate the main workflow; dashed arrows indicate feedback and retraining. B, Overview of fluAgPredictor. A pretrained viral foundation model maps HA and NA sequences into a protein representation space, which is then aligned to HI-derived antigenicity by an attention-based feature extractor. C, Overview of fluVacSelector. Candidate vaccine strains and circulating strains are projected into antigenic space, and pairwise distances are aggregated into a vaccine coverage score for candidate ranking. D, Overview of fluAgEnhancer. Diversity-driven sampling selects representative evolved viruses for HI testing to recalibrate antigenic space through model retraining.

**Figure 2.**
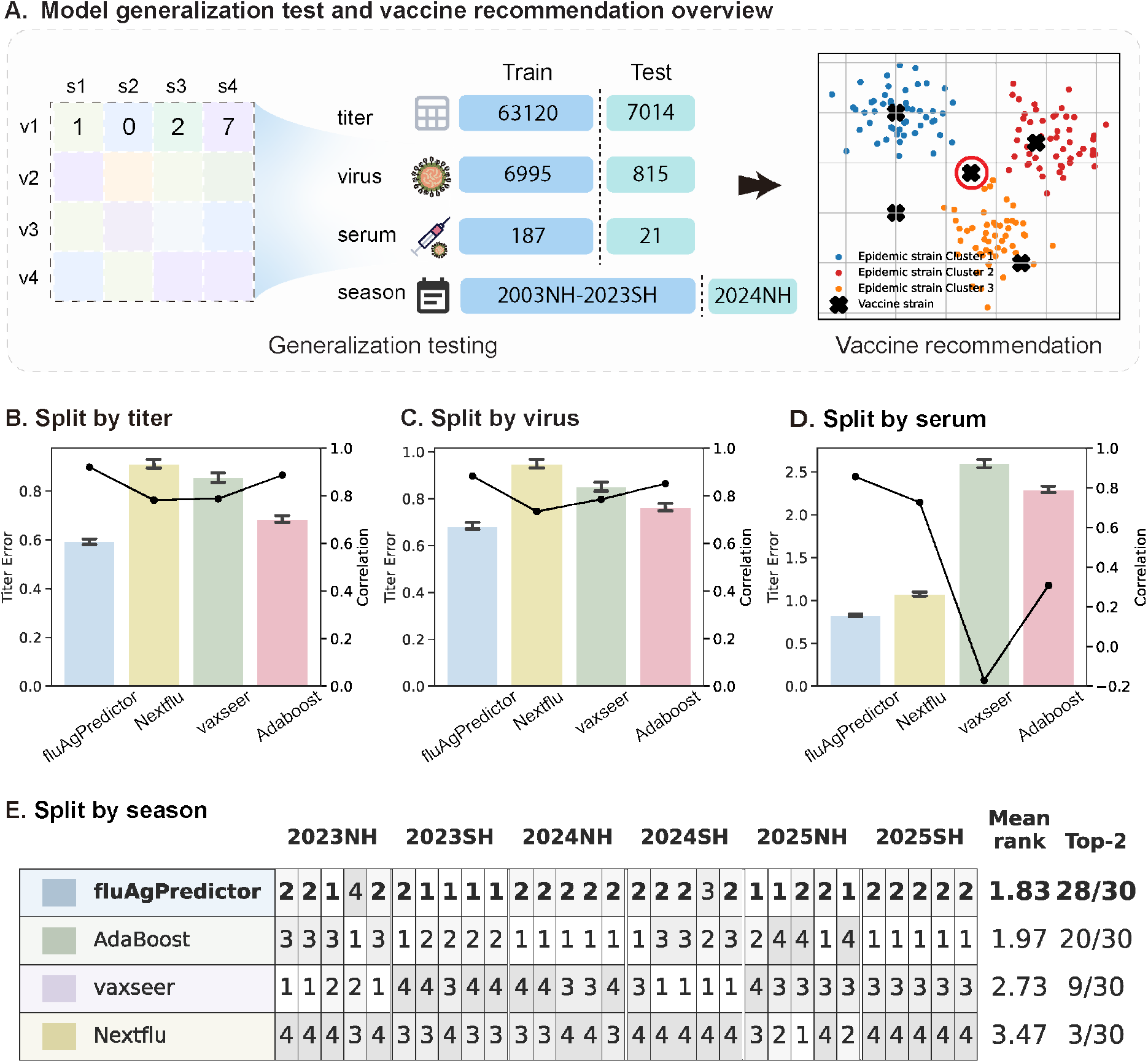
Generalization performance of fluAgPredictor across surveillance-aligned evaluation settings. A, Evaluation design across four surveillance-aligned settings: titer missingness, virus missingness, serum missingness and temporal extrapolation, together with downstream vaccine recommendation. B–D, Performance comparison across models under random removal of 10% titers, 10% viruses or 10% sera, respectively. Bars show error metrics and lines show correlation metrics. E, Rank-based comparison across six held-out influenza seasons from 2023NH to 2025SH. Each season–metric pair was treated as an independent evaluation unit.

After establishing this evidence ladder, the residue-level interpretability of the antigenicity-focused representation was further examined (Fig. 3). Within the same antigenic-space framework, decision-facing utilities were demonstrated, including fluVacSelector (Fig. 1C) for scoring and ranking candidate vaccine strains by predicted coverage and fluAgEnhancer (Fig. 1D) for prioritizing, under limited assay throughput, HI tests on the viruses expected to most improve calibration of the antigenic-space structure, thereby efficiently adding critical phenotypic evidence and enabling continual model refinement.

**Figure 3.**
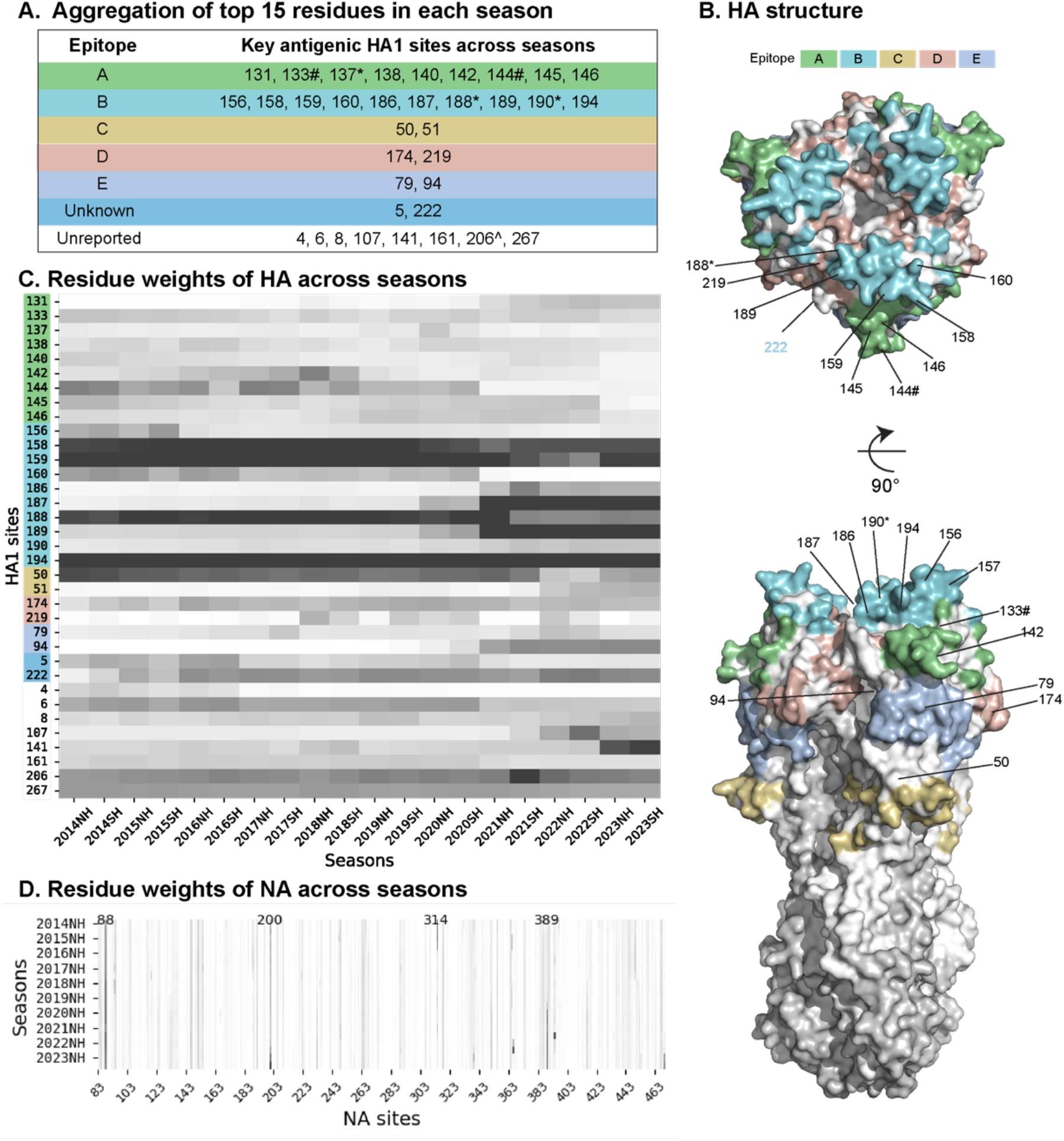
Model-derived antigenic determinants in HA and NA. A, HA1 residues recurrently prioritized across seasons, grouped by classical antigenic epitopes. B, Structural mapping of model-prioritized HA residues on the HA trimer. Two views are shown. C, Heat map of HA residue weights across H3N2 seasons from 2014NH to 2023SH. D, Heat map of NA residue weights across H1N1 seasons.

Together, fluAgPredictor, fluVacSelector and fluAgEnhancer form an integrated workflow that links sequence-based antigenic surveillance with prospective vaccine recommendation, supporting GISRS decision-making under realistic throughput constraints (Fig. 1A).

### Dataset Construction and Model Strategy Optimization

#### Dataset Construction

To develop and evaluate a sequence-based model for predicting influenza antigenic variation, nucleotide and amino acid sequences of HA and NA from H1N1 and H3N2 viruses were integrated with corresponding HI measurements. Sequence data were obtained from GISAID^28-30^, and HI data were compiled from the Francis Crick Institute. HI titers were log2-transformed and converted to antigenic distances by subtracting the homologous titers (see Methods).

The viruses used in the HI assays were matched to their corresponding GISAID entries based on name, passage history, and sampling date, generating paired test-virus, reference-virus, and antigenic distance data.

#### Optimization of Input and Data Strategies

To define a robust baseline under surveillance-realistic missingness and temporal extrapolation, input representations and data strategies were systematically ablated to identify the most stable sequence-to-antigenic-space mapping (Extended Data Fig. 1).

Among the input schemes examined, amino-acid-level representations consistently outperformed nucleotide-based encodings. The joint modeling of NA together with HA, complemented by passage-history metadata, generally further improved predictive accuracy (Extended Data Fig. 1B), consistent with prior observations that NA activity and culture conditions can influence HI readouts. Although subtype-specific training could yield a modest advantage in internal validation, a unified multi-subtype framework showed clearer gains in temporal extrapolation (Extended Data Fig. 1C), supporting the notion that shared structural and evolutionary constraints across lineages can aid forward generalization. Based on these comparisons, the configuration integrating HA/NA embeddings with passage metadata was adopted as the default fluAgPredictor setting for subsequent experiments. Complete ablation results and quantitative comparisons are provided in the Supplementary Information.

#### Foundation Model Selection

Because fluAgPredictor aligns pretrained sequence features to HI-derived antigenicity under sparse supervision, the encoder can materially affect data efficiency and robustness under drift. Therefore, ESM2-3B^12^, LucaOne^31^ and LucaVirus^32^ were compared as sequence encoders while keeping the remainder of fluAgPredictor architecture and training protocol unchanged, and each variant was trained end-to-end with the same antigenic-distance targets (Extended Data Fig. 2).

LucaVirus achieved the strongest overall performance across evaluation settings and was adopted as the default encoder for subsequent experiments (Extended Data Fig. 2). This pattern is consistent with the interpretation that continual pretraining on viral sequence distributions yields embeddings that better capture virus-specific evolutionary and functional constraints, providing a more suitable starting point for antigenicity alignment.

### fluAgPredictor remains robust under sparse and prospective HI evidence

The fluAgPredictor module quantifies antigenic differences between pairs of influenza viruses based on their HA and NA sequences and passage history (Fig. 1A). Increased antigenic distances signify diminished cross-protection between immune responses and serve as pivotal metrics for vaccine potency evaluation and strain selection^26,27,33^.

Robustness was evaluated under four surveillance-aligned scenarios that recapitulate the dominant sources of missingness and the extrapolation faced by the GISRS (Fig. 2A). For a fair comparison, Nextflu, Adaboost, and vaxseer were retrained and validated on the same dataset used for fluAgPredictor, using an identical split protocol (see Methods) to remove confounding from heterogeneous training periods and data coverage.

In the first scenario, partial titer coverage was simulated by randomly masking 10% of entries in the virus–serum HI matrix, reflecting the fact that routine testing does not measure all serum– virus pairs. fluAgPredictor reconstructed the masked values with a mean absolute error (MAE) of 0.593 log2 units and a Pearson correlation coefficient of 0.920, outperforming Nextflu (MAE = 0.911, r = 0.782) and Adaboost (MAE = 0.855, r = 0.788), indicating strong completion of incomplete serological evidence.

In the second scenario, 10% of viruses and their associated titers were randomly withheld during training, evaluating generalization to previously unseen strains. Under this setting, fluAgPredictor retained robust performance (MAE = 0.683, r = 0.882), whereas Nextflu (MAE = 0.949, r = 0.734) and Adaboost (MAE = 0.851, r = 0.785) demonstrated weaker generalization to unseen strains.

The third scenario assessed the generalization to unseen sera by withholding 10% serum columns, reflecting settings in which cross-reactivity must be inferred for newly generated antisera, such as those raised against candidate vaccine strains. fluAgPredictor once again achieved the best performance (MAE = 0.826, r = 0.855), outperforming Nextflu (MAE = 1.074, r = 0.726) and Adaboost (MAE = 2.599, r = -0.170).

Finally, temporal extrapolation was evaluated using six held-out influenza seasons spanning 2023 northern hemisphere (NH) to 2025 southern hemisphere (SH) (2023NH, 2023SH, 2024NH, 2024SH, 2025NH and 2025SH). For each test season, the models were trained exclusively on data from preceding seasons and then evaluated on the target season, thereby mimicking real-world forecasting under strict temporal ordering and preventing information leakage from future seasons. fluAgPredictor was compared with Adaboost, vaxseer and Nextflu using five metrics: MAE, mean squared error (MSE), Pearson correlation, Spearman correlation and R2. To avoid overinterpreting any single season or metric, each season-metric pair was treated as an independent evaluation unit, yielding 30 season-metric units in total. Under this rank-based aggregation, fluAgPredictor ranked within the top two in 28 of 30 units and achieved the best overall average rank (1.83), compared with 20 of 30 units and an average rank of 1.97 for Adaboost. Notably, fluAgPredictor remained in the top two for at least four of five metrics in every test season, whereas Adaboost showed more pronounced season-to-season variability. Across the six seasons, fluAgPredictor also achieved the highest overall Pearson correlation (0.658) and R2 (0.384), while maintaining MAE (0.897), MSE (1.514) and Spearman correlation (0.634) at levels comparable to the best-performing baseline.

Across all four surveillance-aligned scenarios, fluAgPredictor delivered accurate and stable antigenic inference under sparse, incomplete and prospectively accumulated HI evidence (Extended Data Fig. 3). These results are consistent with a more stable sequence-to-antigenic-space mapping under continual drift, with robust generalization across surveillance-relevant missingness and strict temporal extrapolation, supporting the use of fluAgPredictor as a scalable computational complement to experimental HI assays for influenza surveillance and vaccine strain selection.

### Antigenically important sites identified by the model

fluProfiler aims to learn a stable, transferable sequence-to-antigenic-space mapping beyond retrospective fits to pairwise HI readouts. Following evaluation across four surveillance-aligned settings, we examined whether this mapping is supported by mechanistic signal, specifically whether model decisions concentrate on antigenic determinants rather than background sequence variation.

Therefore, residue-level interpretation was conducted in H3N2, a rapidly evolving subtype^34^, to evaluate the stability of model-derived residue importance under continual antigenic drift. Using the task-aligned attention extractor, viral sequences were first mapped to antigenic distances, and the resulting attention weights were then used to derive residue-level importance scores across 20 influenza seasons (2014 NH to 2023 SH). For each season, the 15 residues with the largest attention-derived contributions were identified, and their union defined a cross-season set of key sites (Methods).

Of the significant positions, 27 (77%) mapped to established antigenic epitopes or the sialic-acid receptor-binding site (RBS) (Fig. 3A)^9,35-38^. Sixteen sites clustered in epitopes A and B (Extended Data Fig.4A) and were significantly enriched (one-tailed Fisher’s exact test, P = 1.24 × 10^−5^), recapitulating the established immunodominance of these regions in H3N2 in the absence of any antigenic-site priors^39-41^.

We also analyzed the model’s attention dynamics across the five classical HA antigenic epitopes (A–E) over 20 influenza seasons from 2014 NH to 2023 SH. Epitopes A and B consistently received the highest attention among all regions, with epitope B showing a relative predominance in recent years (Extended Data Fig.4B), consistent with a previous study^39,40^.

At the residue level, a small number of residues exhibited stable prominence across seasons. This stability was concentrated in six residues (158, 159, 188, 194, 206, and 267). Among these, residues 158, 159, 188, and 194 received the highest model attention (Fig. 3C) and clustered on the exposed apex of the HA head within antigenic site B. Residue 188 lies in the 190-helix of the RBS, whereas 158, 159, and 194 map to the binding pocket and influence receptor accessibility. Recurrent substitutions at these sites have been repeatedly associated with antigenic drift and reduced vaccine effectiveness.

In addition to the canonical antigenic hotspots, the model also assigned persistent attention to several less-characterized residues, including HA1-206 and HA1-267, raising the possibility that it captures structural or functional constraints beyond dominant antigenic recognition.

Although HI titers are designed to measure antibody-mediated inhibition of HA binding to sialylated receptors on red blood cells, NA can modulate assay readouts through its receptor-destroying activity by removing terminal sialic acids from red blood cell surface glycoconjugates^42-44^. To capture this effect, NA sequences were incorporated into model training, and attention weights were analyzed to identify key residues (Extended Data Fig.4C). Because oseltamivir is routinely added to H3N2 HI assays to suppress NA activity, attention analyses were restricted to the H1N1 subtype (Fig. 3D). Four high-attention regions were identified at residues 88, 200, 314, and 389. Residue 88 maps to a highly conserved N-linked glycosylation site in N1^45^. Residues 200 and 314 have previously been linked to N1 antigenic evolution^46,47^. Residue 389, in contrast, is less well characterized and has emerged here as an additional candidate site.

Together, these analyses provide a mechanistic interpretation of the learned features of the model, demonstrating their biological plausibility and revealing previously underappreciated HA and NA residues that may shape antigenic evolution within the antigenic space. This integrative approach bridges computational prediction with structural virology, providing a foundation for targeted experimental validation and more efficient influenza surveillance strategies.

### AI-based vaccine strain recommendation

Antigenic characterization is central to seasonal influenza vaccine strain selection^23^. Therefore, we developed fluVacSelector, a prospective framework for ranking candidate vaccine strains according to predicted aggregate antigenic coverage. To mimic the GISRS decision-making process, model training at each consultation was restricted to the antigenic and genetic information available at that time, and this procedure was applied prospectively across six consecutive vaccine composition consultations (Fig. 4A–F).

**Figure 4.**
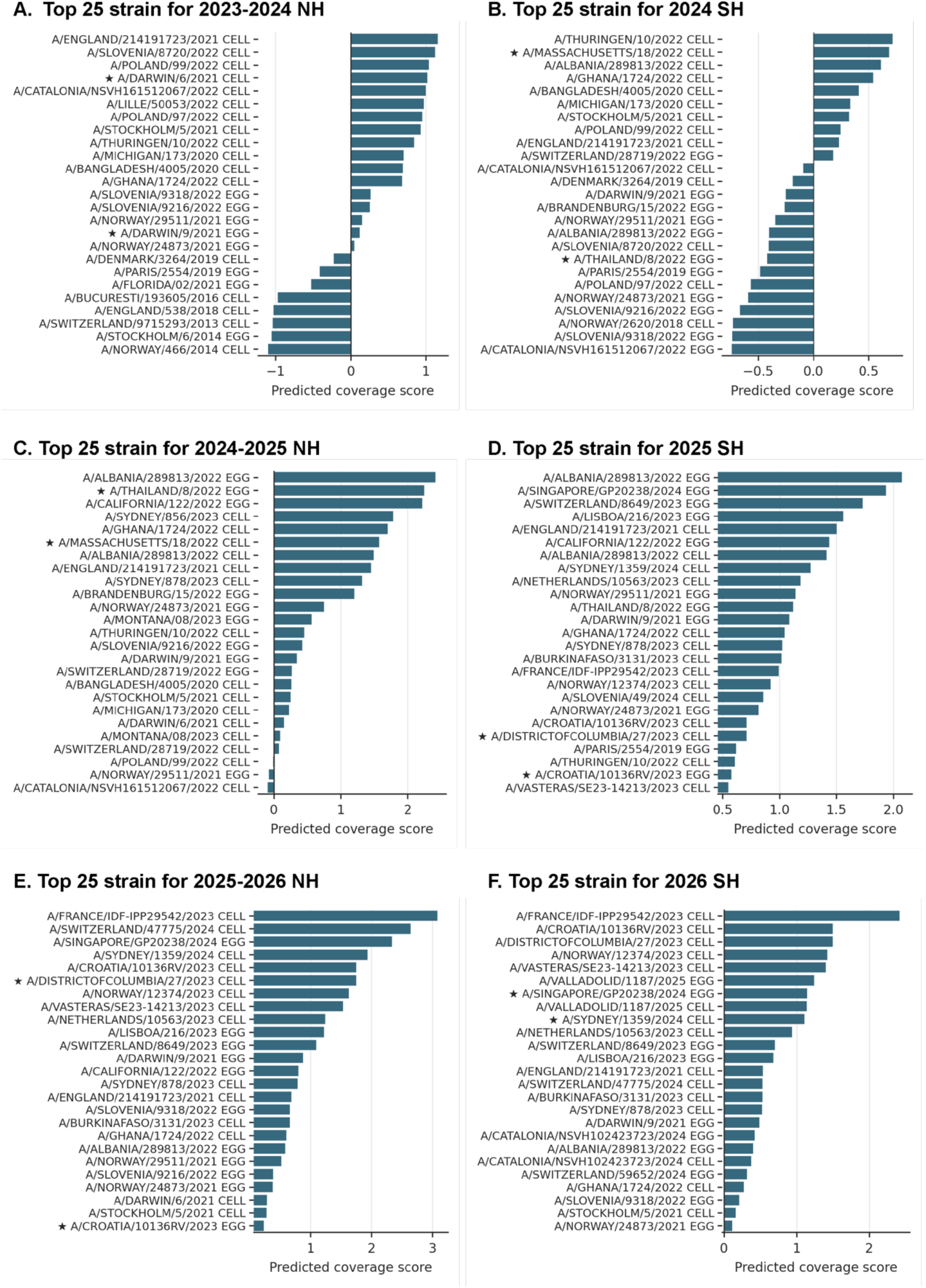
Prospective ranking of candidate vaccine strains by fluVacSelector. A–F, Top 25 candidate strains ranked by predicted aggregate coverage score for six consecutive vaccine composition consultations, spanning 2023–2024 NH to 2026 SH. Asterisks indicate WHO-recommended strains. Higher scores indicate broader predicted antigenic coverage.

Across consultations, WHO-recommended strains consistently ranked within fluVacSelector’s top 25 candidates, indicating that both approaches converged on the same high-coverage region of the candidate space (Fig. 4A–F). At the same time, fluVacSelector also identified higher-scoring alternative candidates in several consultations. Therefore, we examined whether these ranking differences were reflected in the antigenic space and in the underlying sequence changes by comparing fluVacSelector’s top-ranked strains with the passage-matched WHO-recommended strains.

We next projected test viruses and candidate vaccine strains from the Francis Crick Institute into the antigenic space learned by fluProfiler from all data available up to the September 2025 consultation (Extended Data Fig. 5). This projection revealed a clear temporal structure: larger antigenic displacements were usually observed across calendar years, whereas viruses from seasons within the same year generally remained closer in space. In most consultations, the WHO-recommended strain and fluVacSelector-prioritized strain localized near one another and close to the future test-virus cluster. More evident local divergence was observed in the September 2023 and September 2025 consultations, where the WHO-recommended strains also showed larger shifts relative to the previous season. This pattern was consistent with the broader tendency for stronger antigenic displacement across calendar years. Together, these analyses suggest that fluVacSelector generally converges with the overall antigenic direction captured by WHO recommendations, while providing finer discrimination among nearby candidates within the same high-coverage region (Fig. 4A–F; Extended Data Fig. 5).

A site-level comparison provided molecular support for these local differences. Relative to the corresponding passage-matched World Health Organization (WHO)-recommended strains, fluVacSelector’s top-ranked strains differed at residues mapping to known antigenic sites, glycosylation-related motifs, or receptor-binding-site-proximal positions^39-41^ (Extended Data Fig. 6). These changes indicate that the ranking and spatial differences were associated, at least in part, with substitutions in antigenically sensitive regions rather than with background sequence variation alone. These site-level differences place the local ranking differences in a biologically coherent context.

Together, these findings position fluVacSelector as a scalable complement to expert-guided strain selection by extending sequence-guided prioritization across all genomically observed candidates and highlighting additional strains for targeted experimental evaluation (Fig. 4A–F; Extended Data Figs. 5 and 6).

### Diversity-driven active learning supports iterative antigenic model updating under viral evolution

Recent H1N1 and H3N2 viruses followed clear temporal trajectories in a low-dimensional projection of the LucaVirus embedding space, highlighting the continued evolution of these circulating seasonal lineages over time (Fig. 5A, B). Although sequence-based inference provides a degree of extrapolative capacity for newly emerging strains, continued evolutionary accumulation will progressively move circulating viruses beyond the experimental support of historical HI data. Therefore, maintaining an up-to-date antigenic model requires not only accurate sequence-based inference but also efficient prioritization of which viruses to phenotype and incorporate into iterative retraining.

**Figure 5.**
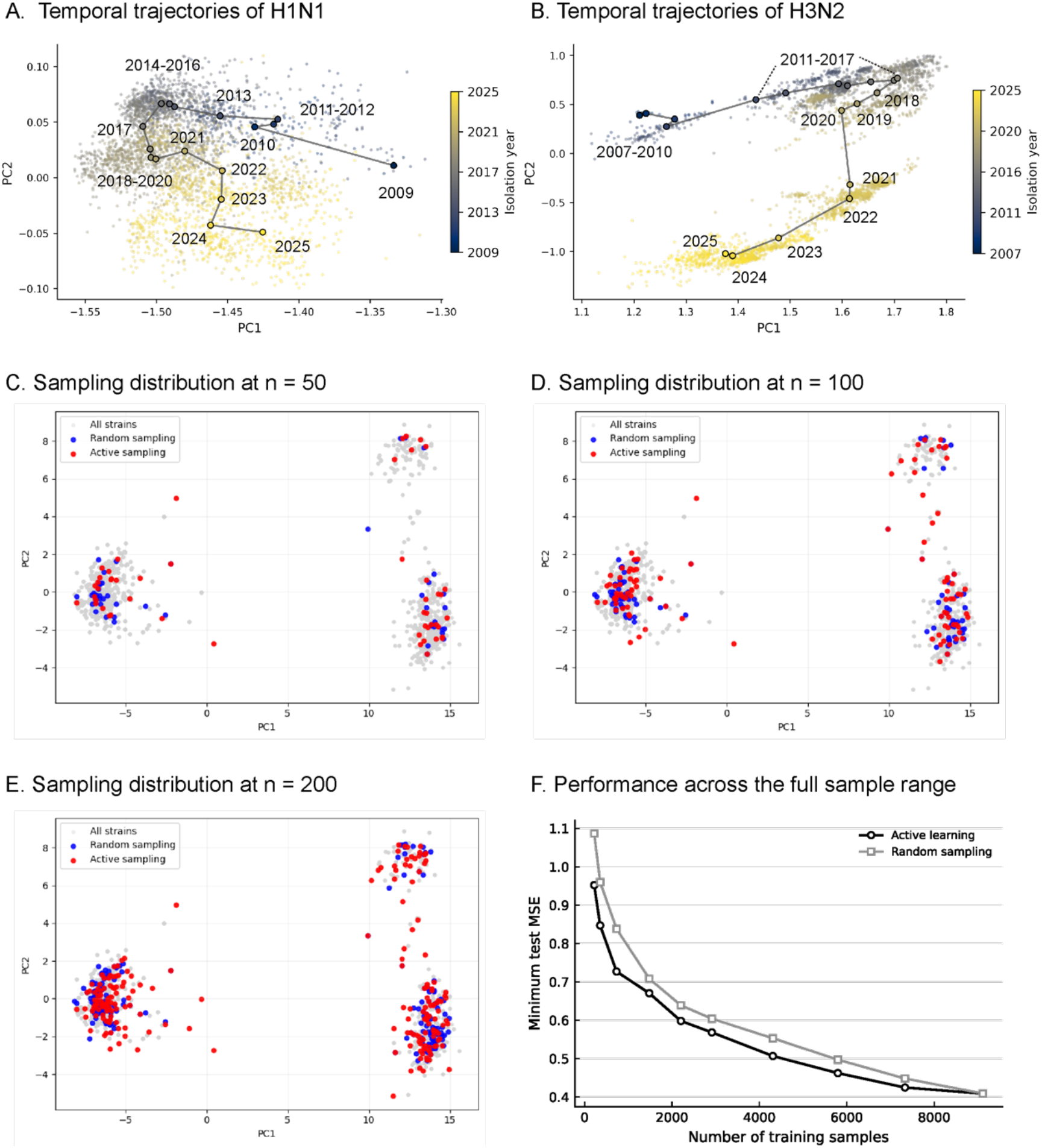
Diversity-driven active learning for iterative antigenic model updating. A,B, Temporal trajectories of H1N1 and H3N2 viruses in a two-dimensional projection of the learned embedding space. Seasonal centroids are connected in chronological order. C–E, Sampling distributions under random sampling and active sampling at budgets of n = 50, 100 and 200, respectively. Grey points indicate all strains, blue points random sampling, and red points active sampling. F, Minimum test MSE across the full sampling range for active learning and random sampling.

The resulting challenge is one of pace: the experimentally anchored antigenic landscape expands more slowly than the diversity of circulating viruses. Because HI assays can be applied to only a limited subset of newly emerging strains in each season, model-refreshing evidence remains sparse and selectively sampled, leaving parts of the evolving antigenic landscape weakly constrained by direct measurement.

To address this challenge, we developed fluAgEnhancer, a data-driven virus selection framework. By optimizing the selection of representative strains for HI testing, fluAgEnhancer improves experimental efficiency, reduces the experimental burden, and expands the model’s exposure to diverse emerging viral patterns.

Across sampling budgets, fluAgEnhancer selected strains that more broadly covered the candidate viral pool in antigenic space while retaining representation of the major circulating clusters (Fig. 5C–E). This broader coverage translated into higher sample efficiency: active learning reached the same predictive target (test MSE=0.5) with approximately 25% fewer training samples and maintained a lower test error across all the sampled ranges (Fig. 5F). Under a fixed experimental throughput, this efficiency gain could instead be used to assay additional viruses, thereby increasing the density of experimentally anchored points and expanding the coverage of the evolving antigenic space.

In summary, these findings show that fluAgEnhancer can stabilize antigenic modeling under ongoing viral evolution when experimental capacity is limited. By coupling sequence-based prediction with model-guided strain selection, it improves the efficiency with which antigenic space can be experimentally anchored and iteratively updated. The gains observed here are likely to be conservative given the scale and coverage of the current dataset. In larger and more diverse surveillance settings, diversity-driven selection may offer even greater value for real-time antigenic surveillance and vaccine strain prioritization.

## Discussion

Within the GISRS, a worldwide laboratory network continuously monitors circulating influenza viruses and generates HI data to support vaccine strain recommendation. Seasonal vaccination remains the most effective defense against influenza, and antigenic characterization based on HI assays is central to this decision-making process. However, extensive sequence diversity and rapid evolution hinder the development of reliable sequence-based models for antigenic prediction and limit the efficiency with which GISRS data can be translated into actionable vaccine recommendations. Here, we present a unified computational framework that operates on the GISRS surveillance system and integrates antigenicity prediction, vaccine recommendations, and data-efficient sampling, with the aim of making antigenic surveillance and vaccine selection more systematic and timelier in a data-driven manner.

Building on the classical view that HI reactivity is primarily governed by the HA1 region, we found that jointly incorporating full-length HA and NA sequences markedly improved predictive accuracy and generalization. This observation does not contradict the paradigm of HA1 immune dominance but instead reveals cooperative HA–NA interactions that are intrinsic to the HI assay. NA provides complementary information via its sialidase activity, glycosylation patterns, and modulation of the HA–NA functional balance, enabling the model to capture antigenic determinants with greater molecular completeness. For the H1N1 datasets generated without the use of NA inhibitors, incorporating NA features effectively captures this complex signal. These results suggest that future HI assays and antigenic modeling efforts should explicitly consider HA– NA cooperativity as a contributor to measured antigenic phenotypes.

In contrast to previous subtype-specific approaches, our multi-subtype, alignment-free strategy improves robustness and generalization across heterogeneous influenza strains. This finding suggests that influenza viruses share conserved antigenic architectures and evolutionary constraints across subtypes, with antigenicity emerging not solely from individual residues—as emphasized in traditional site-specific analyses—but from higher-order combinations of sequence features. As a conceptual complement to conventional mutational studies, we propose that although identifying universal key residues across divergent influenza subtypes remains difficult, transformer-based models can capture shared antigenic signatures in an alignment-free manner, thereby revealing subtype-independent representations of antigenic determinants.

By freezing the parameters of a foundation model and introducing an attention-pooling layer, we sought to balance generalization and interpretability. Attention-weight analyses showed that, in the absence of prior annotation, residues prioritized by the model largely align with known antigenic sites and highlight candidate functional positions (for example, HA1-206 and HA1-267). These findings suggest that deep models may infer structural and biophysical patterns underlying antigenic drift, even without explicit biological priors; however, further experimental validation is required to confirm these predictions and clarify their mechanistic basis.

Owing to the throughput limitations of HI assays, we developed an active-learning sampling strategy that prioritizes antigenically informative and representative viruses within fixed experimental resources. This approach improved predictive accuracy while reducing the number of assays required, thereby minimizing redundant testing and optimizing the allocation of laboratory effort. These results illustrate the potential of AI to enhance experimental design and offer a more efficient framework for the “intelligent surveillance” of rapidly evolving viral populations.

Several limitations of this study should be noted. First, the current model treats NA features similarly across datasets, regardless of whether NA inhibitors were used in the HI assays. Explicitly modeling such assay-condition heterogeneity using richer experimental metadata could further enhance robustness and calibration. Second, this study focuses on seasonal H1N1 and H3N2 because these subtypes provide large, standardized HI datasets for modeling. In contrast, influenza B and zoonotic lineages remain sparsely characterized, which may limit generalization across circulating influenza diversity. As more data accumulate, extending the framework to these lineages should provide a more comprehensive global antigenic landscape. Third, HI measurements are subject to biological and technical variability (including ferret immune heterogeneity and passage-associated changes), which may bias antigenicity estimates. Incorporating additional phenotypes (e.g., microneutralization) and sequencing the exact isolates used in HI assays could further improve robustness.

Efforts can further improve this work. The framework can realize its greatest value when embedded in a global surveillance system as a bidirectional, continual-learning loop. Rather than merely ingesting GISRS data, a mature system should feed operational guidance back into the network, prioritizing viruses that fall in sparsely covered regions of the antigenic landscape inferred from sequence representations. Each new round of phenotyping then adds targeted sequence– phenotype pairs, progressively sharpening antigenic resolution and keeping the model calibrated under ongoing viral evolution.

With continual recalibration and refinement of the sequence–antigenicity mapping, we can perform in silico mutagenesis based on currently circulating strains and evaluate the antigenic consequences of these simulated variants. Building on efforts to predict viral evolution, we can leverage the learned sequence–antigenicity mapping to design and pre-screen scalable libraries of virtual antigens with broad coverage against future strains (e.g., peptide and mosaic vaccine constructs). Ultimately, this could help drive a paradigm shift in vaccine strain selection, transforming it from a largely reactive process into a proactive, continually self-improving framework.

Currently, we have focused this framework on influenza vaccine decision-making, given the relatively rich landscape of sequencing and antigenic surveillance data. As surveillance infrastructures mature and datasets accumulate, we anticipate that this “copilot” paradigm can be extended across a diverse range of viral and bacterial pathogens. More broadly, our aim is not to build yet another predictor tailored to a single pathogen, but to establish a general, operational methodology that is both data-efficient and transferable—one that converts routine surveillance into actionable, forward-looking immunological risk assessment to support proactive management of emerging threats.

## Supporting information

Extended Documentation

## Methods

### Dataset source and curation

HA and NA sequence data of human seasonal influenza A viruses were retrieved from GISAID, including both nucleotide and corresponding translated amino acid sequences for H1, H3, N1, and N2 entries submitted between January 1, 1975, and March 31, 2025. Sequence records from human seasonal H1N1 and H3N2 viruses were retained for downstream analysis. Amino acid sequences were used in the final fluAgPredictor framework and downstream analyses, whereas nucleotide sequences were included for comparative evaluation of alternative input representations during model strategy optimization.

HI data were compiled from 44 surveillance reports released by the Francis Crick Institute and covering influenza seasons from 2003 to 2025SH. These reports provided HI measurements for circulating/test viruses against post-infection ferret antisera raised to reference viruses. In total, 115,927 HI measurements were collected, comprising 59,920 measurements for H1N1 and 56,007 for H3N2. For each virus isolate, we extracted available metadata from the reports, including the virus name, passage history, and collection date.

Sequence and antigenic data were matched at the virus-isolate level. For each isolate, HA and NA sequences were paired with the corresponding HI record using strain identity and associated metadata. Passage histories were manually standardized into two assay-relevant categories, egg and cell, and the combination of strain name and passage category was used to define a unique virus isolate throughout the study. Virus-antiserum pairs with invalid HI measurements or ambiguous passage categories were excluded. Records that could not be reliably matched to the required sequence entries and metadata were also removed. The resulting curated dataset was used for model development, evaluation and downstream analyses.

### Construction of HI-derived antigenic distance

For each virus-antiserum measurement, the HI reactivity was converted into a continuous HI-derived antigenic distance using a normalized HI titer (NHT)-based formulation. Let *T*_*aβ*_ denote the measured HI titer of test virus a against an antiserum *β*, where *β* was raised against a reference virus b. Let *T*_*bβ*_ denote the corresponding homologous HI titer of the reference virus *b* against its matched antiserum *β*. We then defined the NHT-based antigenic difference for the virus-antiserum pair as:

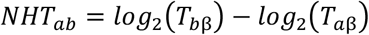

This transformation quantifies the reduction in cross-reactivity of the test virus relative to the homologous reference and expresses all measurements on a common log2 scale. Larger values indicate greater antigenic divergence between the test and reference viruses recognized by the antiserum, whereas a value of zero indicates reactivity equivalent to the homologous reference measurement. The resulting NHT-based antigenic difference was used as the regression target throughout model development, evaluation and downstream analyses.

Because antigenic measurements are defined with respect to a specific antiserum, the same virus can yield different target values when evaluated against antisera raised to different reference viruses. This formulation preserves the assay-defined asymmetry of HI measurements while enabling direct learning from paired sequence and serological data.

### Surveillance-aligned data splits and evaluation metrics

To evaluate model performance under surveillance-relevant deployment settings, we designed four complementary data-splitting protocols that reflect distinct sources of incompleteness encountered in routine influenza antigenic surveillance: missing virus-antiserum measurements, previously unseen viruses, previously unseen antisera, and future seasonal drift. All models were trained and evaluated using identical data partitions to ensure fair comparison. In the titer-removal setting, individual virus-antiserum measurements were randomly partitioned into training, validation and test sets at a ratio of 8:1:1. This setting evaluates matrix completion performance, that is, the ability to infer missing antigenic relationships for virus-antiserum pairs, both of whose component viruses and antisera had both been observed during training.

In the virus-removal setting, virus isolates were first partitioned into training and test groups at a ratio of 9:1 at the isolate level, and all measurements involving test viruses were withheld from the model fitting. Measurements associated with the remaining training viruses were then randomly divided into training and validation subsets at a ratio of 8:1. This setting evaluates the generalization to previously unseen viruses.

In the serum-removal setting, antisera were partitioned analogously such that all measurements involving held-out antisera were assigned to the test set, and measurements involving the remaining antisera were further randomly split into training and validation subsets at a ratio of 8:1. This setting evaluates generalization to previously unseen antisera.

To assess prospective generalization under ongoing viral evolution, we further performed strict temporal extrapolation across six held-out influenza seasons: 2023 NH, 2023 SH, 2024 NH, 2024 SH, 2025 NH and 2025 SH. For each test season, the models were trained exclusively on data from all preceding seasons, with no access to HI measurements, virus isolates or antisera from the target season. A validation subset was reserved from the historical pre-test data for model selection. This design prevents temporal information leakage and mimics the real-world forecasting scenario in which antigenic relationships must be inferred for newly emerging viruses using only previously available data.

Model performance was quantified using five complementary metrics: mean absolute error (MAE), mean squared error (MSE), Pearson correlation coefficient, Spearman correlation coefficient, and coefficient of determination (R2). For the six-season temporal extrapolation analysis, each season metric combination was treated as an independent evaluation unit, resulting in 30 units in total. Within each unit, models were ranked from best to worst according to the corresponding metric, with lower values indicating better performance for MAE and MSE and higher values indicating better performance for the Pearson correlation, Spearman correlation and R2. The overall performance across seasons was then summarized using the average rank across all 30 units, together with the number of units in which a model ranked within the top two. This rank-based aggregation was used to assess both predictive accuracy and cross-season stability without overemphasizing any single season or metric.

For all comparative experiments, the same evaluation framework was applied to fluAgPredictor and all baseline models.

### Modeling strategy optimization and foundation-model comparison

To define a robust default configuration for subsequent analyses, we systematically evaluated alternative input representations, antigen-component combinations, data strategies and pretrained sequence encoders under the surveillance-aligned benchmarking framework described above. These analyses were designed to identify a practical modeling setup that remained accurate under realistic missingness and temporal extrapolation, rather than to maximize performance in any single evaluation scenario.

To evaluate the modeling strategy, we compared nucleotide-based and amino-acid-based sequence representations, different antigen component combinations, and alternative metadata settings. We assessed models using HA alone, NA alone, or combined HA and NA inputs, with or without passage history information. These comparisons were performed using a shared downstream prediction architecture so that performance differences primarily reflected the contribution of the input and data strategy rather than changes in model capacity. Across evaluation settings, amino acid representations consistently outperformed nucleotide-based representations, and the combined use of HA, NA, and passage-history information provided the most stable overall performance. Therefore, this configuration was adopted as the default input strategy for subsequent model development.

Next, we compared multiple pretrained sequence foundation models as upstream encoders using the same downstream architecture and evaluation protocol. This analysis was intended to isolate the effect of representation quality while keeping the prediction head and benchmarking setting fixed. Among the tested encoders, LucaVirus achieved the strongest and most consistent overall performance across the surveillance-aligned evaluation settings. Therefore, LucaVirus was selected as the default sequence encoder in the final fluAgPredictor framework.

### fluAgPredictor architecture and training

fluAgPredictor was designed to estimate the HI-derived antigenic distance between a reference and a test virus directly from their HA and NA protein sequences, along with passage-history information. The final model architecture consisted of three components: a sequence foundation model, a task-specific feature extractor, and a regression head.

For each virus pair, the HA and NA amino acid sequences of the reference and the test viruses were separately encoded by the selected foundation model to generate residue-level embeddings. Based on the comparative analyses described above, LucaVirus was adopted as the default sequence encoder for all subsequent experiments. The passage history for the two viruses was represented as categorical metadata indicating egg or cell passage and was encoded using a dedicated passage encoder.

The sequence branch processes four protein inputs per sample: HA of the reference virus, NA of the reference virus, HA of the test virus, and NA of the test virus. For each input sequence, residue-level embeddings produced by the foundation model are passed to an attention-pooling layer, which assigns task-specific weights to different positions and aggregates the sequence into a fixed-length representation. This design allows the model to retain information from the pre-trained viral representations while emphasizing the most informative residues for antigenic discrimination. In parallel, passage metadata from the two viruses are embedded into a learnable vector representation through the passage encoder. The four sequence-derived representations and the passage-derived representation are then concatenated and passed to a multilayer perceptron (MLP) regression head to predict a single continuous output corresponding to the NHT-based antigenic distance.

The model hyperparameters were kept fixed across all experiments. The hidden dimension of the sequence encoder output was 2560, the attention-pooling dimension was 256, and the passage encoder produced a 256-dimensional representation. The concatenated feature vector was processed by a projection block with two fully connected hidden layers having 512 and 128 units, respectively, using ReLU activation and a dropout rate of 0.1, followed by a final linear layer for the regression output.

All fluAgPredictor models were trained using the AdamW optimizer with a learning rate of 1 × 10 - 4 and a batch size of eight. The training objective was the MSE loss between the predicted and observed NHT-based antigenic distances. Training and inference were implemented in PyTorch 2.2.1 and performed on an NVIDIA A100 80 GB GPU with CUDA 12.1. Unless otherwise specified, the same training setup was used for all comparative experiments.

### Baseline models and fair retraining protocol

To ensure a fair comparison across methods, all baseline models, including Nextflu, Adaboost and vaxseer, were retrained and evaluated on the same curated HI-sequence dataset used for fluAgPredictor in the titer-removal, virus-removal, serum-removal and temporal extrapolation benchmarks, rather than compared using previously reported results obtained from different datasets, surveillance periods and evaluation protocols.

For each evaluation setting, the baseline models inherited the same training, validation and test partitions used for fluAgPredictor. In the titer-removal setting, all models were trained on the same 80% of observed virus-antiserum measurements, tuned on the same validation subset and evaluated on the same held-out 10% test subset. In the virus- and serum-removal settings, the same held-out virus isolates or antisera were used across all methods. In the temporal extrapolation analysis, all methods were trained exclusively on data from seasons preceding the target season and were evaluated on the same held-out season, with model selection performed using only pre-test historical data. This unified protocol ensured that any performance differences reflected differences in modeling capacity rather than differences in data access.

Nextflu was implemented as a representative classical antigenic prediction framework that models pairwise antigenic differences from sequence-derived genetic information, along with virus- and serum-associated effects. Adaboost was implemented as a non-linear ensemble regression baseline trained on the same paired antigenic-distance data. In addition, vaxseer was evaluated as a recent deep-learning baseline. Each baseline retained its own modeling framework; however, all were retrained under the same data partitions and evaluated using the same target definition and performance metrics as fluAgPredictor.

Model performance for all methods was quantified using the same five metrics: MAE, MSE, Pearson correlation coefficient, Spearman correlation coefficient, and coefficient of determination (R2). For temporal extrapolation across six held-out seasons, rank-based aggregation across the 30 season-metric evaluation units was applied identically to fluAgPredictor and all baseline models.

### Residue-level interpretation, epitope aggregation, and enrichment statistics

To assess whether the learned sequence-to-antigenic-space mapping captured biologically meaningful determinants of antigenic variation, we performed residue-level interpretability analyses using the task-aligned attention extractor of fluAgPredictor. Because the foundation model provides general contextual sequence representations whereas antigenicity-specific reweighting is introduced by the downstream attention-based extractor, the interpretation focused on the learned attention patterns of the trained predictor rather than on the pre-trained embeddings alone. For each evaluated virus pair, token-level attention scores were extracted from the task-specific feature extractor and converted into residue-level importance profiles by aggregating attention weights over the feature dimension. The residue-level scores were then averaged across evaluation pairs to obtain season-level importance estimates.

Primary residue-level interpretation was conducted for H3N2, a rapidly evolving subtype with sustained antigenic drift across recent surveillance seasons. We analyzed 20 influenza seasons spanning 2014-2023. For each season, residues were ranked by their season-averaged importance scores, and the top 15 positions were retained as the most influential sites for that season. The union of these seasonal top-ranked positions defined the cross-season set of key HA sites used in downstream analyses. The season recurrence of each site was also recorded to identify residues showing stable prominence across multiple seasons.

To evaluate biological concordance, selected HA positions were mapped to curated annotations of the five classical H3 antigenic epitopes (A–E) and to residues associated with the receptor-binding site (RBS). The enrichment of key sites within epitopes A and B was assessed using a one-tailed Fisher’s exact test, comparing the observed number of selected positions in these epitopes with the number expected from the full set of analyzed HA positions. In parallel, season-level attention scores were aggregated across residues belonging to each of the five classical epitopes to quantify epitope-level attention dynamics over time. These aggregated scores were used to compare the relative contribution of epitopes A–E across the 20 analyzed seasons.

To distinguish antigenically exposed from potentially structural sites, recurrent high-attention residues were further examined in the context of sequence conservation and structural location. Residues that repeatedly appeared among high-ranking positions across seasons were classified as season-stable candidates and subsequently interpreted with reference to known epitope topology, RBS proximity and solvent exposure in the HA head region.

Because neuraminidase (NA) can indirectly influence HI readouts through modulation of red-blood-cell receptor cleavage, we also analyzed attention-derived importance patterns in NA. However, because oseltamivir is routinely added to H3N2 HI assays to suppress NA activity, NA-focused interpretability analyses were restricted to the H1N1 subtype. Residue-level attention scores were derived using the same procedure as for HA, and season-aggregated profiles were used to identify high-attention regions in the N1 protein for downstream biological interpretation.

### Prospective vaccine candidate ranking with fluVacSelector

fluVacSelector is a deterministic downstream ranking module built on top of fluAgPredictor to prioritize candidate vaccines with broad predicted antigenic coverage for a given consultation setting. For each recommendation exercise, all input data were truncated to those available before the corresponding consultation date, including HI measurements, sequence records and passage metadata, thereby mimicking the prospective decision context of routine influenza surveillance.

For a predefined set of candidate vaccine strains and a circulating virus panel, fluVacSelector used fluAgPredictor to estimate pairwise HI-derived antigenic distances *d*(*c*_*i*_, *v*_*j*_) between each candidate strain ci and each virus vj in the reference panel from HA and NA sequences, along with passage information. Egg- and cell-propagated versions of the same strain were treated as distinct candidates because passage-associated substitutions may alter antigenic phenotypes and therefore affect predicted coverage.

For each candidate vaccine strain *c*_*i*_, we computed an aggregate mismatch score across the reference panel as follows:

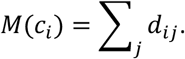

where smaller values indicate lower overall predicted antigenic mismatch to the reference panel. For visualization and ranking, this quantity was converted into a predicted performance score:

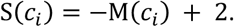

The additive constant of 2 anchors the score to a commonly used reference point in HI-based antigenic analysis, where 2 antigenic units correspond to a fourfold HI difference. This shift was introduced for interpretability only and does not affect candidate ranking. Candidate vaccine strains were therefore ranked in descending order of *S*(*c*_*i*_), and the top-ranked candidates were interpreted as those with the most favorable predicted overall match to the viruses represented in the reference panel.

This procedure was applied prospectively to each consultation-specific recommendation task using only contemporaneously available data. In this study, fluVacSelector was used to generate candidate rankings for the 2023-2024 NH to 2026 SH vaccine recommendations. The module introduced no additional trainable parameters and functioned as a deterministic post-processing layer on top of the trained fluAgPredictor model.

### Structural annotation of vaccine-difference sites

To interpret the molecular features underlying differences between model-prioritized and consultation-selected vaccine strains, we performed post hoc structural annotation of vaccine-difference sites for each prospective recommendation setting. For each consultation, the number of amino acid differences between the top-ranked candidate identified by fluVacSelector and the corresponding WHO-recommended vaccine strain was enumerated and analyzed at the residue level.

Differential residues were mapped to curated annotations of canonical H3 antigenic epitopes, residues proximal to the receptor-binding site (RBS), and predicted N-linked glycosylation motifs. Potential N-linked glycosylation sites were defined as N-X-S/T motifs, where X denotes any residue except proline. Differences were further interpreted in the context of exposed loop regions surrounding the HA head and RBS-adjacent structural elements to assess whether candidate-specific substitutions were likely to affect epitope accessibility, glycan shielding or local receptor-binding geometry.

This analysis was used as a mechanistic interpretation layer for prospective vaccine ranking and did not contribute to the fluVacSelector scoring procedure itself. Instead, it provided a biologically informed explanation of how sequence differences between candidate strains may reshape antigenically exposed surfaces and thereby contribute to differences in predicted coverage.

### Diversity-driven sampling for model updating with fluAgEnhancer

fluAgEnhancer is a diversity-driven active-learning module designed to prioritize viruses for HI phenotyping and model updating under limited experimental throughput. Rather than selecting newly emerging viruses at random, fluAgEnhancer identifies a subset of representative viruses that more broadly spans the evolving sequence-derived antigenic landscape, thereby improving the efficiency with which new phenotypic evidence is added to the model.

For each candidate virus, the HA amino-acid sequence was first encoded by the same sequence foundation model used in fluAgPredictor to generate residue-level embeddings. These embeddings were then processed by a trained task-specific feature extractor, which applied learned position-specific attention weighting to produce a pooled sequence representation. The pooled representation was further transformed through a linear projection layer to obtain a fixed-length 256-dimensional task-aligned embedding for each candidate virus. Applying this procedure to all candidate viruses yielded an embedding set that served as the representation space for diversity-driven sampling.

To select a representative subset under a predefined sampling budget k, candidate-virus embeddings were partitioned into k clusters using k-means clustering under Euclidean distance. For each cluster, the virus with an embedding closest to the cluster centroid was selected as the representative strain for HI testing. This procedure generated a subset of viruses that preserved broad coverage of the candidate pool while reducing redundancy from densely populated regions of the representation space.

Selected viruses were then added to the training set for model updating, and the resulting model was evaluated on a held-out test set to quantify the gain in predictive performance achieved under each sampling budget. As a non-adaptive baseline, random sampling was performed under the same budget constraints and evaluation procedure. Comparative performance was assessed using the test MSE after model updating, and sampling efficiency was summarized by the number of selected viruses required to reach a predefined predictive target.

This framework was used to evaluate whether model-guided diversity sampling could improve antigenic model updating under ongoing viral evolution. Because fluAgEnhancer operates on the learned sequence-to-antigenic-space representation rather than on raw sequence identity alone, it preferentially selects viruses expected to add complementary phenotypic information for recalibrating the antigenic-space structure.

## Data availability

The HA and NA protein sequences and associated metadata used in this study were obtained from GISAID (https://gisaid.org/). The accession identifiers for all sequences analyzed in this study, together with the processed matched datasets and source data underlying the main and extended figures and tables, are provided in the project repository (https://github.com/Chengxugorilla/fluProfiler) and/or Supplementary Data. HI data were compiled from the annual and interim reports of the Worldwide Influenza Center at the Francis Crick Institute (https://www.crick.ac.uk/research/platforms-and-facilities/worldwide-influenza-centre/annual-and-interim-reports). Information on human influenza vaccine compositions was obtained from the GISAID resource portal (https://gisaid.org/resources/human-influenza-vaccine-composition/). All other data supporting the findings of this study are available in the manuscript, its Supplementary Information, or from the corresponding author upon reasonable request.

## Code availability

The code used for data processing, model training, evaluation and figure generation in this study is publicly available in fluProfiler GitHub repository (https://github.com/Chengxugorilla/fluProfiler). The repository also contains the scripts required to reproduce the main benchmarking analyses, vaccine-ranking analysis and diversity-driven sampling experiments.

## Author contributions

Y.S. and Z.L. conceptualized and supervised the research. Y.C., Y.X. and Yanhui Cheng prepared and curated the data. D.W. provided the HI data resources. Y.C. developed the fluProfiler framework. Y.C. evaluated the performance of fluProfiler models and comparative machine-learning methods. Y.C. performed the molecular analyses. T.B., J.Y., H.L. and L.Z. contributed to the interpretation of the results. X.Q., X.D., L.Y. and M.S. contributed to the design of the validation strategy. Y.C., Y.S. and Z.L. interpreted the results. Y.C. wrote the original draft of the manuscript. Y.S. and Z.L. critically revised the manuscript. All authors read and approved the final manuscript.

## Acknowledgements

We thank the Chinese National Influenza Center for support with HI data collection and curation. We also thank the Worldwide Influenza Centre at the Francis Crick Institute for sharing antigenic data.

## Competing interests

The authors declare no competing interests.

**Extended Data Fig. 1.**
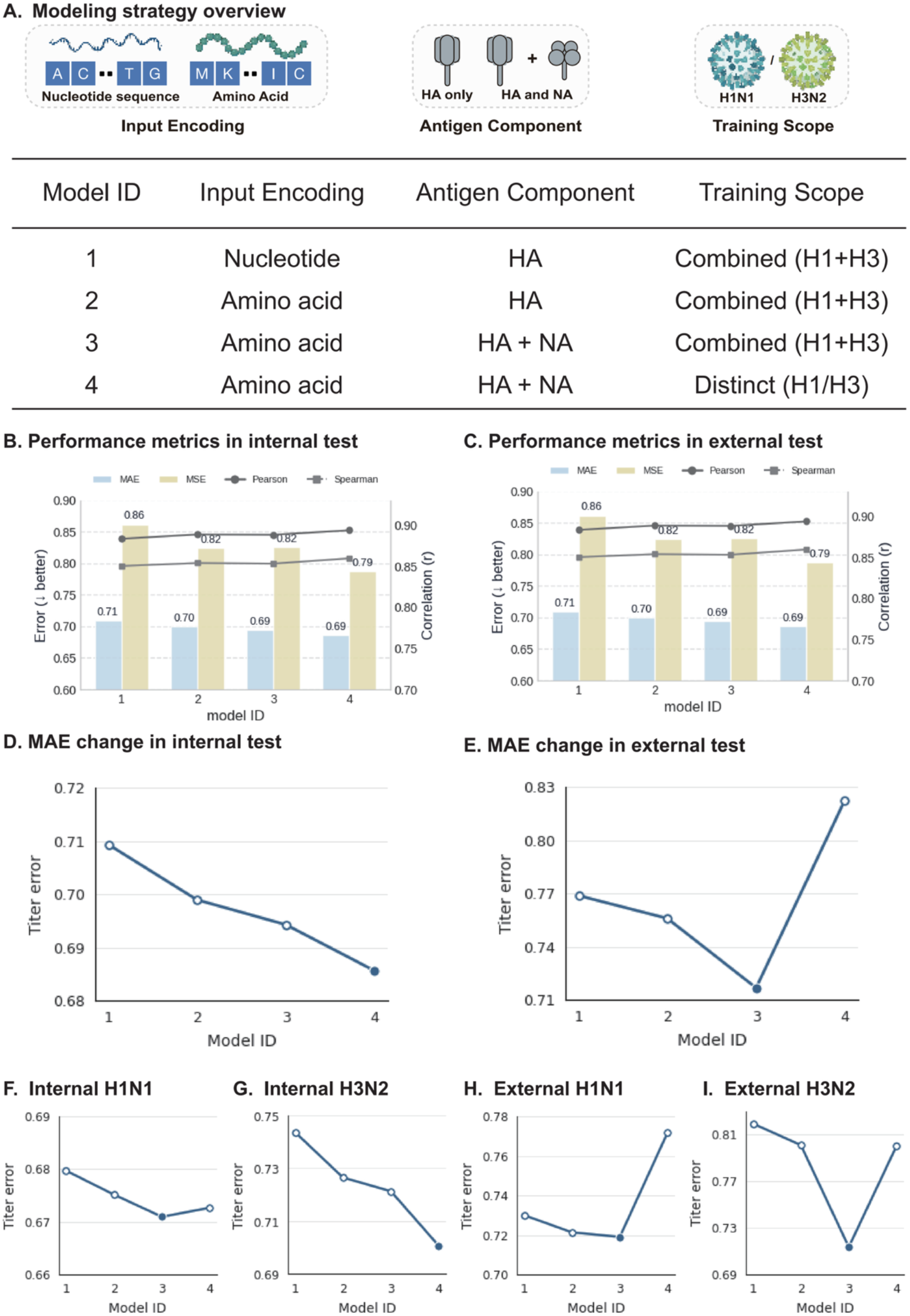
Optimization of input representation and training strategy. A, Overview of the compared modeling strategies. B,C, Performance metrics across strategies in internal and external test sets, respectively. D,E, MAE changes across strategies in internal and external test sets, respectively. F–I, Subtype-specific comparisons in H1N1 and H3N2.

**Extended Data Fig. 2.**
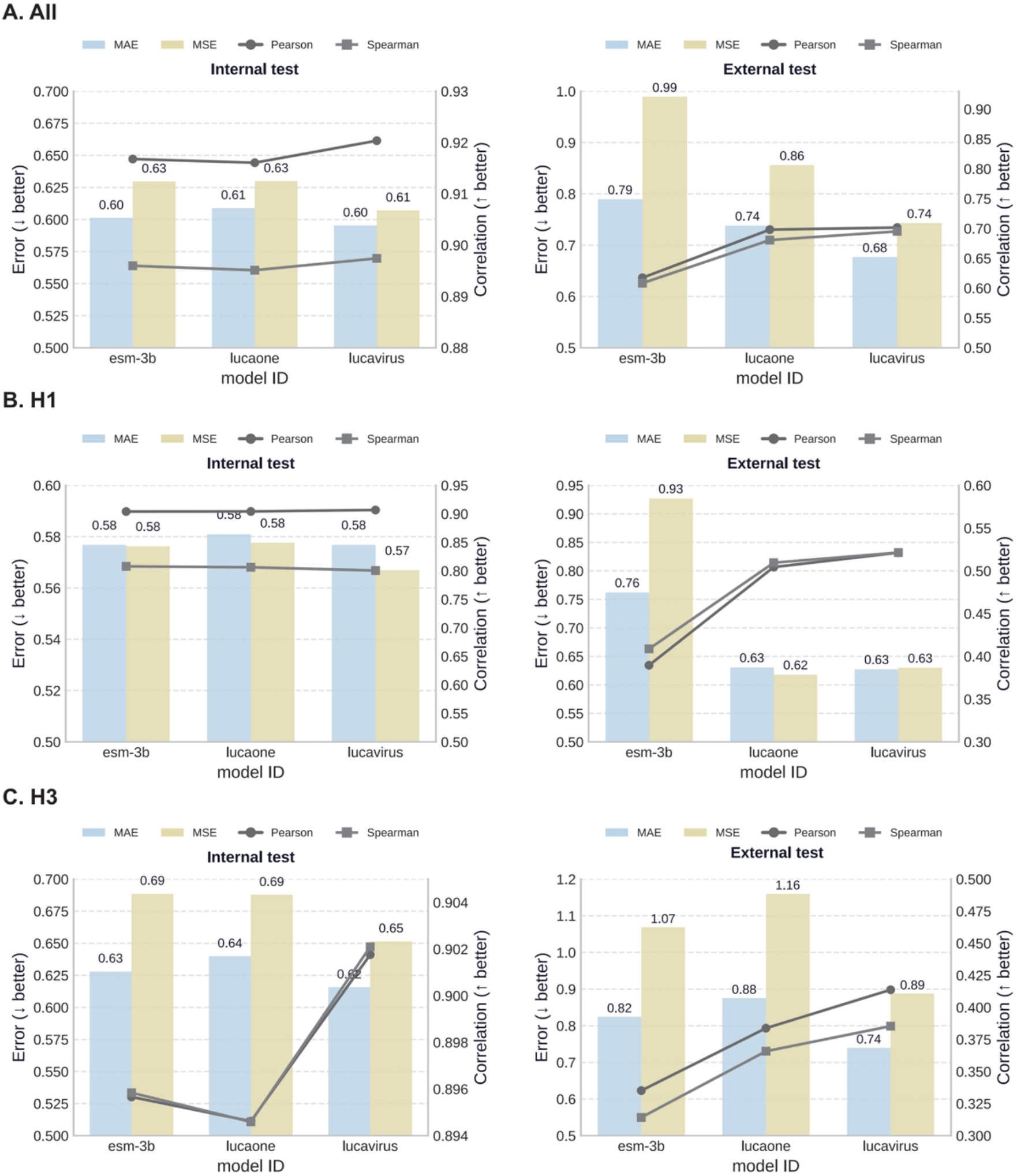
Comparison of alternative foundation-model encoders. A, Encoder comparison on the full dataset. B,C, Encoder comparison in H1N1 and H3N2, respectively. Internal and external test results are shown for each encoder.

**Extended Data Fig. 3.**
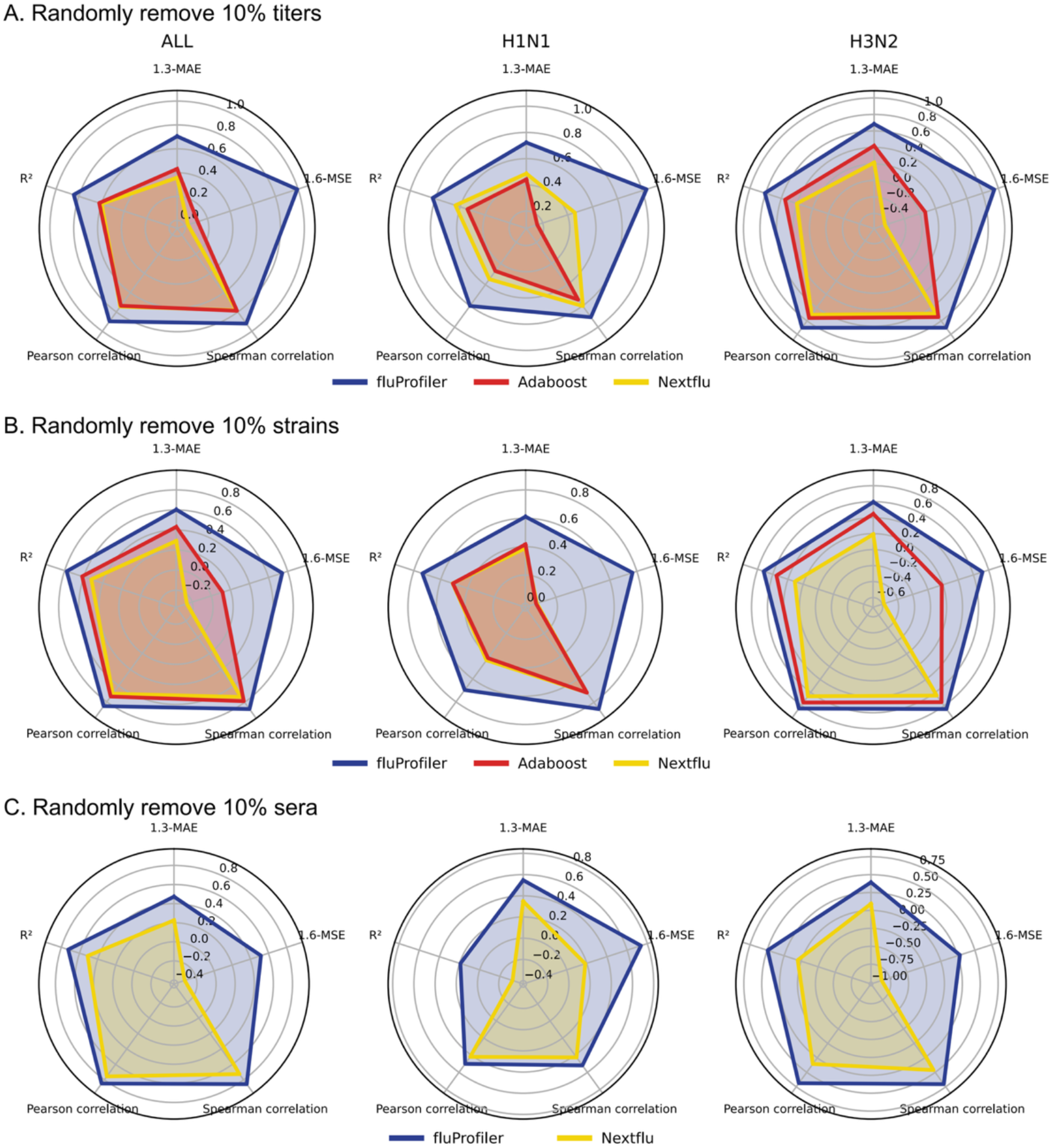
Comprehensive comparison under three missingness settings. A–C, Radar plots summarizing model performance after random removal of 10% titers, 10% viruses or 10% sera, respectively, shown for all strains, H1N1 and H3N2.

**Extended Data Fig. 4.**
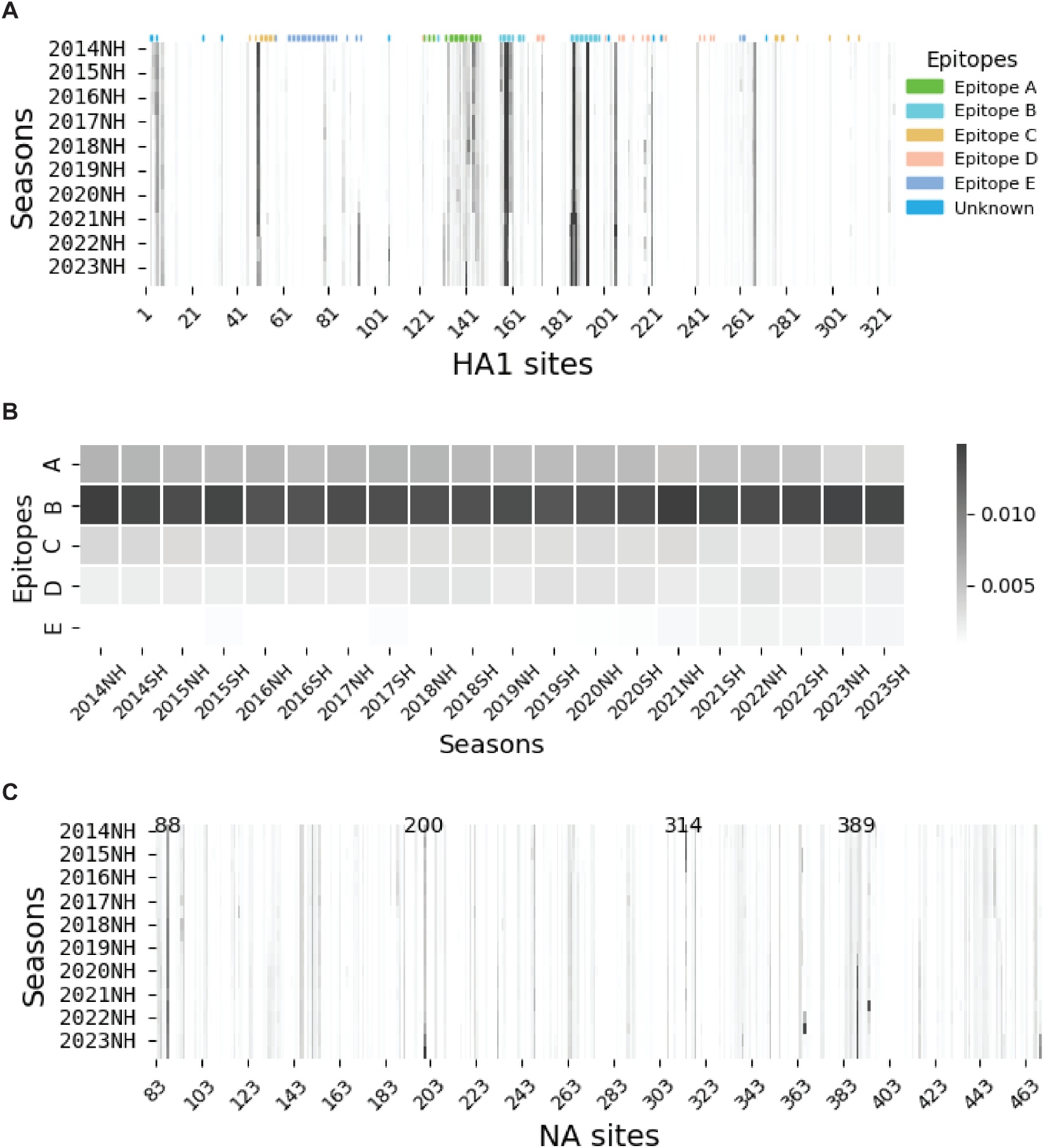
Additional analyses of model-derived residue importance. A, Seasonal heat map of HA1 residue attention scores in H3N2. B, Epitope-level attention weights across seasons. C, Seasonal heat map of NA residue attention scores in H1N1.

**Extended Data Fig. 5.**
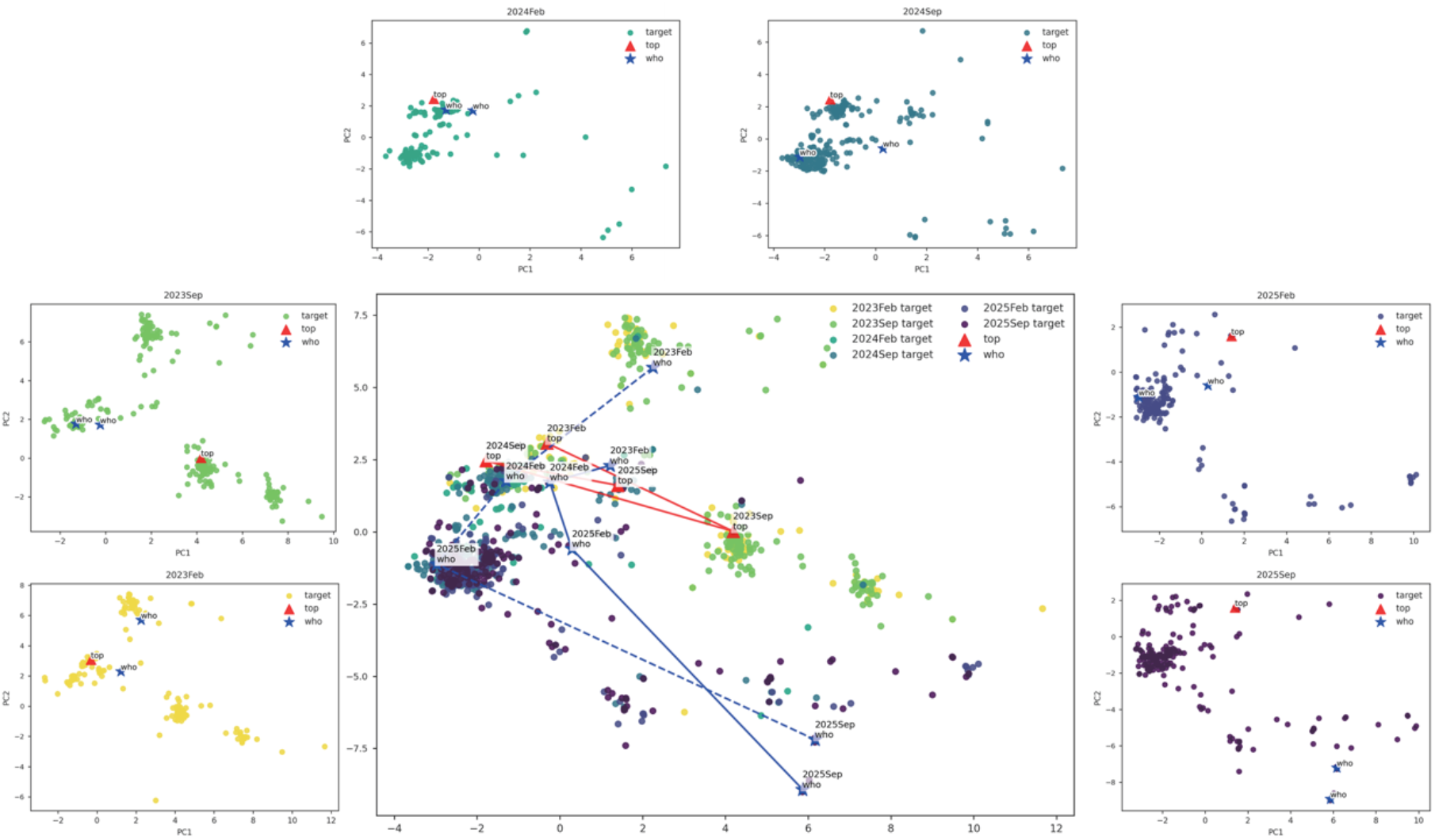
Antigenic-space visualization of candidate ranking across consultation windows. Two-dimensional projections of the learned antigenic space showing target viruses together with the fluVacSelector top-ranked strain and the WHO-recommended strain for each consultation. Insets show individual consultations; the center panel summarizes their relative trajectories in the shared space.

**Extended Data Fig. 6.**
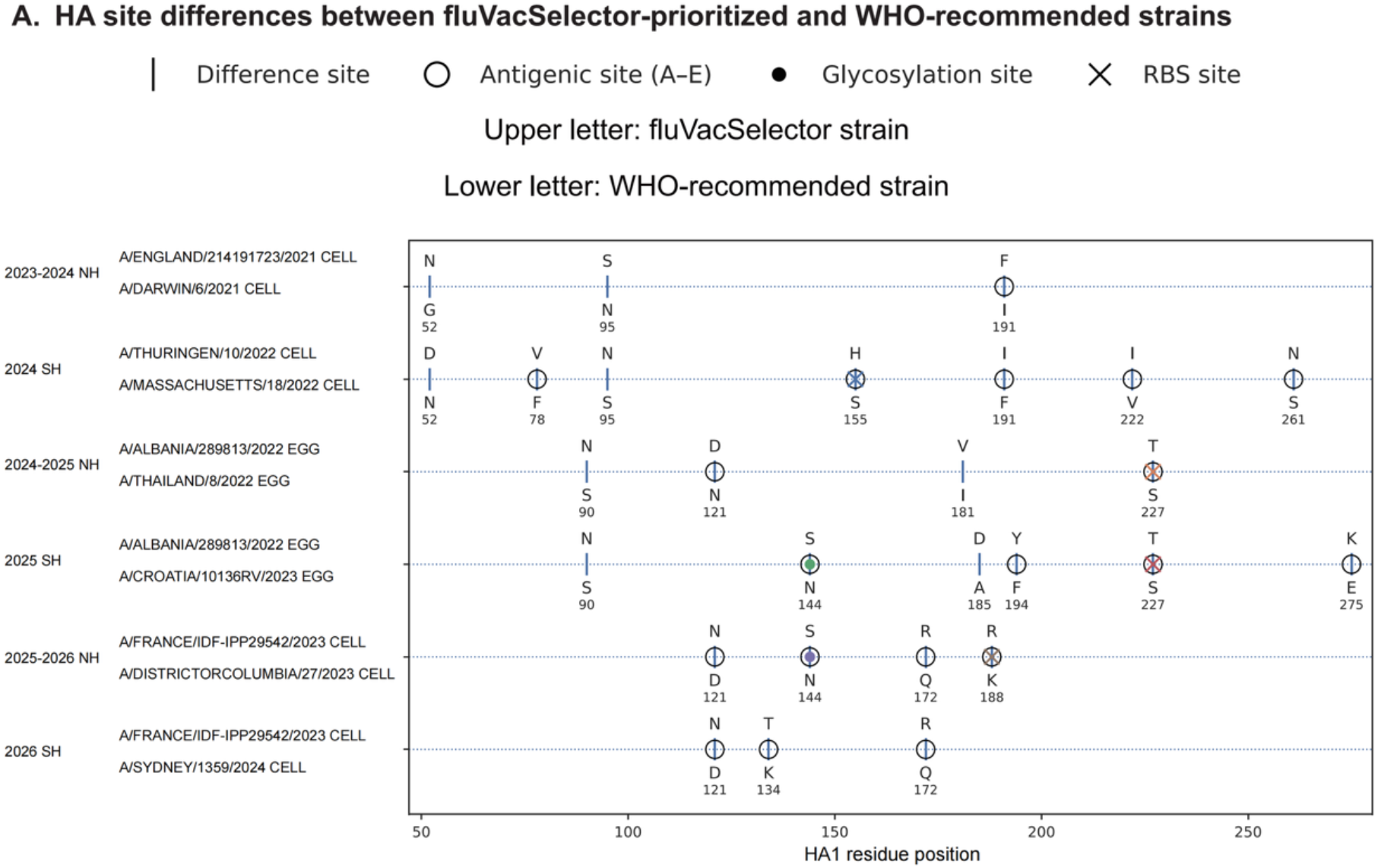
HA site differences between fluVacSelector-prioritized and WHO-recommended strains. Amino-acid differences between the top-ranked fluVacSelector strain and the corresponding WHO-recommended strain across six vaccine composition consultations from 2023-2024 NH to 2026 SH. Upper letters indicate the fluVacSelector strain and lower letters the WHO-recommended strain. Symbols denote antigenic sites, glycosylation sites and receptor-binding-site positions.

**Extended Data Fig. 7.**
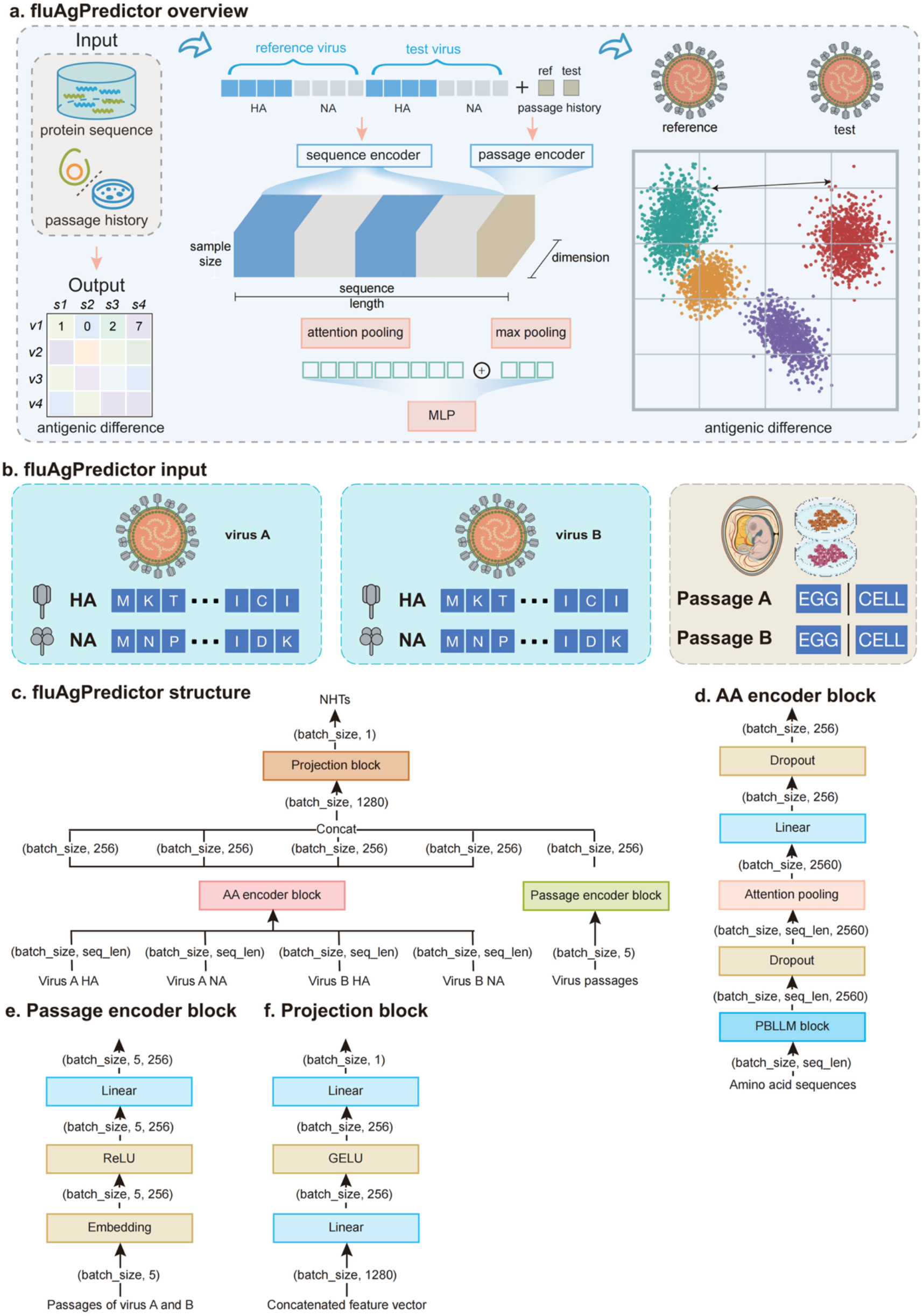
Detailed architecture of fluAgPredictor. A, End-to-end overview of fluAgPredictor. B, Input representation of paired viruses and passage metadata. C, Overall module composition. D, Amino-acid encoder block. E, Passage encoder block. F, Projection block.

**Extended Data Fig. 8.**
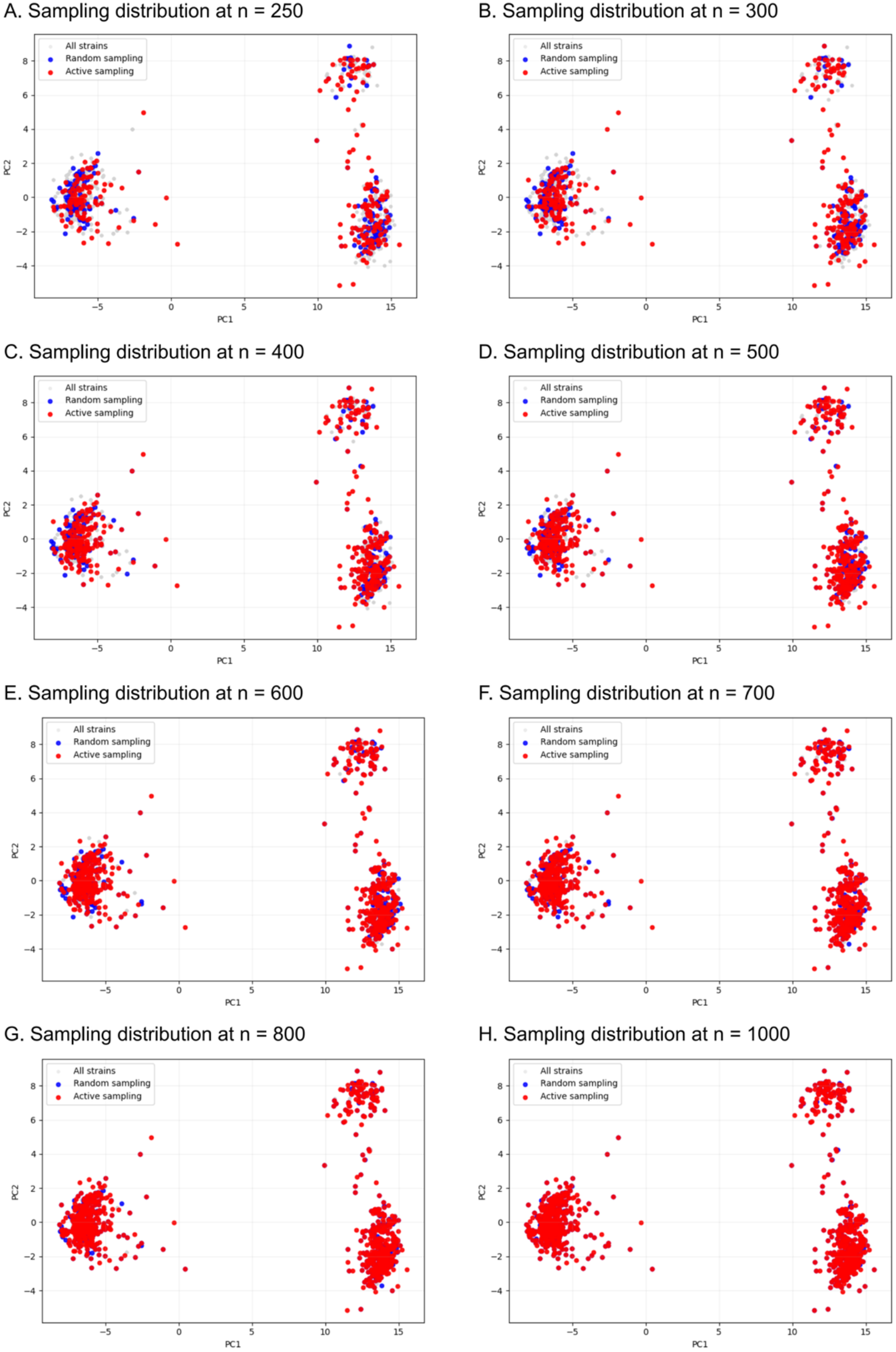
Sampling distributions across larger active-learning budgets. A–H, Sampling distributions at budgets of n = 250, 300, 400, 500, 600, 700, 800 and 1,000, respectively. Grey points indicate all strains, blue points random sampling, and red points active sampling.

