## Extended Documentation for "A sequence-based proactive intelligence for influenza antigenic profiling improves vaccine strain selection"

**Optimization of Input and Data Strategies**

The fluAgPredictor module uses viral hemagglutinin (HA) and neuraminidase (NA) protein sequences together with passage-history metadata as model inputs. This design was motivated by both assay biology and the need for a stable sequence-to-antigenic-space mapping. HA is the major determinant of influenza antigenic variation and the principal driver of hemagglutination inhibition (HI) readouts. Although NA does not mediate hemagglutination directly, its sialidase activity can influence the extent of agglutination and thereby modulate measured HI titers in an assay-dependent manner. In addition, viruses with identical amino-acid sequences may still display different antigenic phenotypes depending on passage history, potentially reflecting differences in host-cell adaptation and post-translational processing.

To define a robust default configuration under surveillance-realistic missingness and temporal extrapolation, we systematically benchmarked alternative input representations and data strategies within a common lightweight Moving Average Equipped Gated Attention (MAGA) architecture with fixed model capacity, so that performance differences could be attributed to input or training design rather than architectural variation (Extended Data Fig. 1).

Across the input schemes examined, amino-acid-level representation consistently outperformed nucleotide encoding, reducing prediction error in both internal and external (seasonal split) data (MAE 0.709 to 0.698 and MAE 0.768 to 0.755, respectively; Extended Data Fig. 1B, C). Within the same architecture, amino-acid inputs also improved computational efficiency by shortening the effective sequence length.

Incorporating NA sequences alongside HA further improved accuracy (internal MAE 0.694 versus HA-only; external MAE, 0.716 versus 0.755; Extended Data Fig. 1B), indicating that NA-derived sequence information can provide complementary signal for HI-related antigenic inference. Notably, for H3N2 HI datasets, neuraminidase inhibitors (for example, oseltamivir) are routinely used to suppress NA activity, enabling reliable agglutination and minimizing NA-driven confounding. Under such assay conditions, NA sequence features may contribute less to the observed HI signal and could act as nuisance variation. This assay-dependent effect likely contributes to subtype- and dataset-specific differences in the benefit of including NA.

Although subtype-specific models yielded a slight advantage in internal validation, unified multi-subtype training improved temporal extrapolation performance (held-out MAE, 0.716 versus 0.880 for subtype-specific training; Extended Data Fig. 1C), suggesting that shared structural and evolutionary constraints across subtypes support more stable forward generalization.

Based on these comparisons, the configuration integrating HA and NA sequence features with passage-history metadata was adopted as the default fluAgPredictor setting for subsequent experiments. Detailed quantitative comparisons are provided in Extended Data Fig. 1 and the Supplementary Information.
